# Rapid RPA-CRISPR/Cas12a detection of *Vibrio coralliilyticus* for coral aquaculture disease surveillance

**DOI:** 10.64898/2026.10.15.759329

**Authors:** Sean Asgari, Jason Doyle, Clare M. Grimm, David G. Bourne, Yui Sato

## Abstract

Coral aquaculture is expanding globally to support both the ornamental aquarium trade and reef-restoration initiatives, yet disease outbreaks remain a threat to production, causing potential substantial coral mortality and economic losses. Effective disease management requires early pathogen detection; however, conventional molecular diagnostics are labor-intensive and poorly suited to time-sensitive aquaculture operations. Here, we developed a rapid diagnostic assay for the coral pathogen *Vibrio coralliilyticus* by implementing recombinase polymerase amplification (RPA) with Clustered Regularly Interspaced Short Palindromic Repeats (CRISPR)/Cas12a-based fluorescent signaling for sensitive and specific detection. The assay was optimized and validated against DNA from seven *V. coralliilyticus* strains and seven non-target *Vibrio* species, demonstrating high specificity and a detection limit <10 target copies per reaction within 30 minutes following DNA extraction. Performance of the assay was further evaluated using water from flow-through aquaria containing healthy and diseased *Acropora tersa* colonies, and results independently validated using droplet digital PCR (ddPCR). Although RPA-CRISPR/Cas12a consistently detected *V. coralliilyticus* across the tanks holding both healthy and diseased corals, ddPCR indicated substantially higher pathogen abundance in diseased tanks, suggesting that pathogen load rather than the presence alone is a more reliable indicator of disease risk. Incorporating rapid semi-quantification of pathogen abundance in future assay development could therefore improve disease prediction and management. This study demonstrates the potential of CRISPR-based diagnostics as a fast, sensitive, and field deployable tool for routine coral aquaculture biosecurity, enabling earlier intervention and more effective mitigation of disease outbreaks.

**Importance:** Disease outbreaks threaten coral aquaculture operations that support reef restoration and the ornamental aquarium trade, highlighting the need for rapid and reliable detection to monitor disease risks in aquaculture systems. However, conventional detection methods are often slow and labor-intensive, limiting their suitability for routine surveillance. Here, we developed a rapid molecular diagnostic method that detects *Vibrio coralliilyticus*, a recognized coral pathogen, within 30 minutes of sample processing. The method successfully detected the pathogen in water collected from coral broodstock tanks, demonstrating its potential application in coral aquaculture. By substantially reducing detection time, this approach offers a model assay development pipeline as a practical foundation for more accessible and timely pathogen surveillance. This work advances molecular diagnostics for marine pathogens and provides a potential tool to strengthen disease monitoring and inform health management of marine organisms in captivity, supporting aquaculture biosecurity and the production for marine restoration and the aquarium trade.

## Introduction

Driven by rapid expansion of reef restoration initiatives and shifts within the global ornamental aquarium trade toward cultivated stocks (Rhyne et al., 2012; Suggett et al., 2023; Wood et al., 2012), demand for cultured corals has increased. Land-based (*ex situ)* aquaculture provides a sustainable production platform while enabling environmental conditions to be carefully controlled to maximize coral growth and survival (Barton et al., 2017; Knapp et al., 2022; Leal et al., 2016). However, intensive aquaculture systems also create conditions that can increase physiological stress, facilitate pathogen transmission, and promote disease outbreaks, particularly when corals are maintained at high densities within interconnected water systems (Cain, 2022; Sheridan et al., 2013; Sweet et al., 2012). Such outbreaks can spread rapidly through production facilities, causing extensive coral mortality and significant economic losses (Sheridan et al., 2013).

Coral diseases commonly manifest as tissue loss, discoloration, growth anomalies, or characteristic banding lesions, yet these visual signs often appear only after infections are well established (Moriarty et al., 2020; Richardson, 1998). Consequently, visual inspections alone rarely provide sufficient warning for timely interventions. Disease processes are often complex and multifactorial, arising through interactions between environmental stressors, opportunistic microorganisms, and the coral host, rather than a single causative agent (Bourne et al., 2009; Thurber et al., 2025; Voolstra et al., 2024). Coral health is governed by the stability of the coral holobiont, a dynamic partnership between the animal host, Symbiodiniaceae, bacteria, archaea, fungi, viruses, and other microorganisms (Bourne et al., 2009). Environmental stress can destabilize this association, promoting microbial dysbiosis and increasing the host’s susceptibility to opportunistic and pathogenic microbes (Gignoux-Wolfsohn et al., 2017). Among bacterial pathogens, members of the genus *Vibrio* are the best characterized (Munn, 2015; Rosenberg & Ben-Haim, 2002), with *Vibrio coralliilyticus* recognized as a principal bacterial pathogen of reef-building corals owing to its well-defined virulence mechanisms and repeated associations with disease outbreaks and coral mortality (Ben-Haim et al., 2003; Sussman et al., 2008; Ushijima et al., 2014, 2020).

Early pathogen detection is essential for effective disease management in aquaculture systems. Quantitative polymerase chain reaction (qPCR) remains the benchmark for pathogen detection and quantification (Bustin, 2017; Rieder et al., 2025), including for *V. coralliilyticus* (Chimetto Tonon et al., 2017; Pollock et al., 2010). More recently, droplet digital PCR (ddPCR) has emerged as a highly sensitive alternative that enables absolute quantification without requiring external standard curves (Pinheiro et al., 2012) and has been applied to coral aquaculture surveillance (Grimm et al., 2026; Ushijima et al., 2020). Despite their analytical strengths, both qPCR and ddPCR require specialized infrastructure, expensive instrumentation, and multi-hour workflows limiting their utility for rapid decision-making applications in aquaculture facilities (Bustin, 2017; Hindson et al., 2011; Tan et al., 2023).

Isothermal amplification technologies address many of these limitations by amplifying nucleic acids at a constant temperature, eliminating the need for thermocycling equipment while substantially reducing assay time (Boonbanjong et al., 2022; Notomi et al., 2000; Oliveira et al., 2021; Piepenburg et al., 2006). Among available isothermal amplification methods, recombinase polymerase amplification (RPA) is particularly attractive because it requires only a single primer pair, simplifying assay development, and typically generates detectable amplicons in 10-20 min (Lobato & O’Sullivan, 2018; Oliveira et al., 2021). Clustered Regularly Interspaced Short Palindromic Repeats (CRISPR)-associated (Cas) proteins have transformed molecular diagnostics through programmable recognition of specific nucleic acid sequences (Kaminski et al., 2021). Cas12a is a CRISPR RNA (crRNA)-guided endonuclease that, following target recognition, exhibits collateral cleavage activity against single-stranded DNA (ssDNA) reporters, generating a fluorescent signal for highly sensitive detection (Chen et al., 2018; Paul & Montoya, 2020). CRISPR/Cas12a platforms have been successfully applied to a wide range of microbial pathogens, including several *Vibrio* species, demonstrating rapid, sensitive, and highly specific detection suitable for point-of-need diagnostics (Li et al., 2022; Wang et al., 2024; Xiao et al., 2021; Zhong et al., 2025). Combining RPA with CRISPR/Cas-based detection provides both rapid amplification and exceptional analytical specificity (Kaminski et al., 2021), while its use has not been evaluated in aquaculture settings for corals.

Here, we present a proof-of-concept RPA-CRISPR/Cas12a assay for the rapid and specific detection of *V. coralliilyticus*, focusing on Indo-Pacific strains relevant to Great Barrier Reef coral aquaculture. We systematically optimized primer sets, crRNAs, and reaction conditions to maximize analytical sensitivity and specificity before validating the assay using water samples collected from aquaria holding healthy and diseased corals. By providing pathogen detection in under 30 minutes following DNA extraction, this study establishes the feasibility of RPA-CRISPR diagnostics for routine coral aquaculture surveillance and lays the foundation for rapid, operational molecular tools that will improve biosecurity and enable earlier intervention to support coral health monitoring and disease management.

## Materials and Methods

### Bacterial culturing and DNA purification

Seven strains of *V. coralliilyticus* and seven non-target *Vibrio* species were used to evaluate assay specificity (Table 1). Isolates were cultured on Marine Agar 2216 (BD Difco, USA) overnight at 28 °C before single colonies were inoculated into 3 mL Marine Broth 2216 (BD Difco) and incubated overnight at 28 °C at 180 rpm. Bacterial cells were harvested by centrifugation (4561 x g, 10 min, 20 °C; Allegra V-15R with VS 4.750 Swinging-Bucket Rotor; Beckman Coulter, USA) and genomic DNA extracted with the DNeasy Blood and Tissue Kit (Qiagen, Germany) according to the manufacturer’s instructions. DNA concentration and purity were determined using a NanoDrop 2000 spectrophotometer (Thermo Scientific, USA). Species identity was confirmed by full-length 16S rRNA gene amplification followed by Sanger sequencing (Macrogen, South Korea). Resulting sequences were compared against the NCBI nucleotide database using BLASTN to verify identification of each isolate (Benson et al., 2013).

**Table 1.**
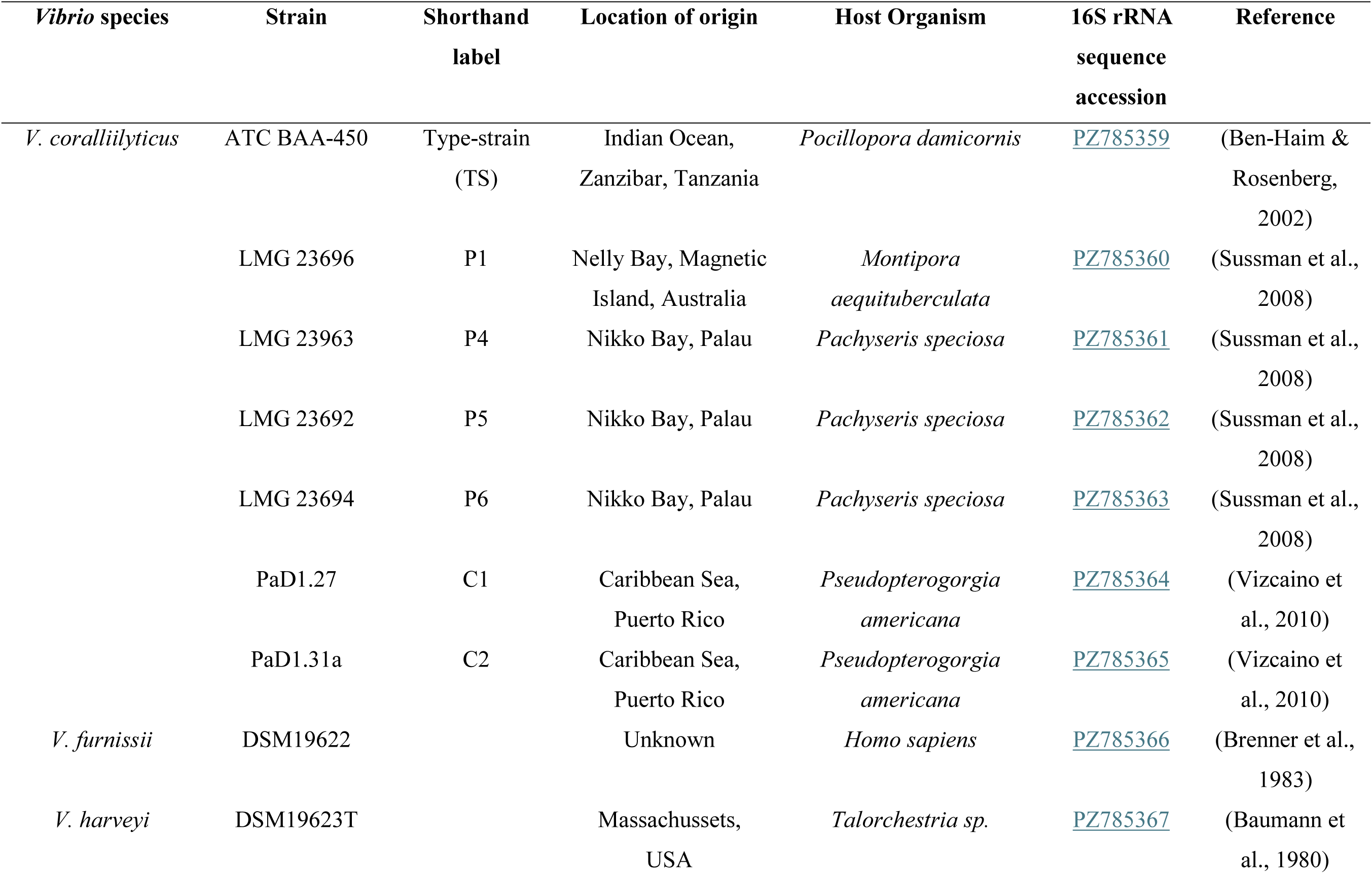
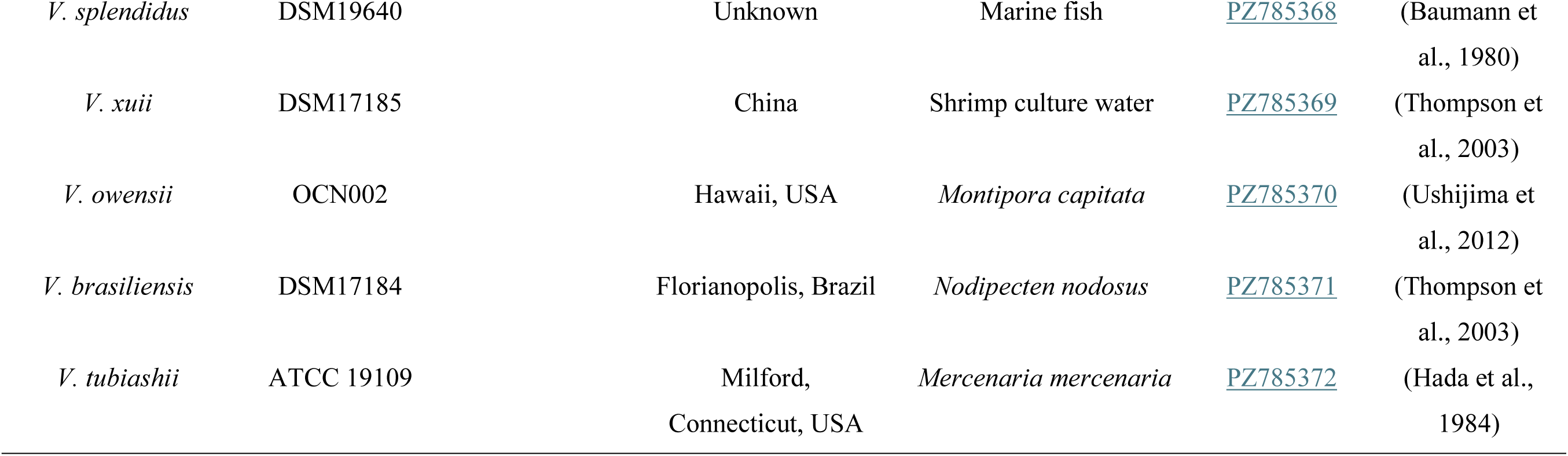
*Vibrio* species and strains used in the development of the RPA-CRISPR/Cas12a assay for the detection of *V. coralliilyticus*.

### ddPCR assay optimization

To establish a quantitative foundation, ddPCR was performed using the Bio-Rad QX200™ ddPCR System (Bio-Rad, USA) with EvaGreen Supermix for absolute quantification of *V. coralliilyticus* DNA. Primers targeting the d*naJ* (heat shock protein 40) gene of *V. coralliilyticus* developed by Pollock et al. (2010) were used throughout. Assay conditions were initially optimized using genomic DNA from the *V. coralliilyticus* type-strain by evaluating primer concentrations ranging from 200 to 700 nM and annealing temperatures from 52 to 66 °C. Optimal conditions were selected based on clear separation of positive and negative droplet populations, consistent amplification, and minimal background fluorescence. These conditions were subsequently validated using five additional *V. coralliilyticus* strains (excluding strain C2) and seven non-target *Vibrio* species (Table 1). All ddPCR reactions were performed in sealed 96 well plates with two technical replicates for each primer concentration and annealing temperature combinations.

Droplet fluorescence data were analyzed using QuantaSoft™ Analysis Pro (Bio-Rad, USA) with manual thresholding. Assay performance was evaluated using the Z-prime factor (Z’; Equation 1) as per Zhang et al. (1999), signal-to-noise ratio (SNR; Equation 2), percent coefficient of amplitude variation (%CV; Equation 3) for both positive and negative droplet populations, and absolute DNA concentration (copies µL^−1^).

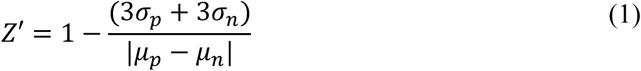

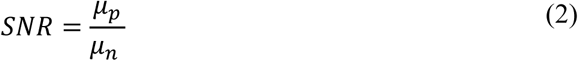

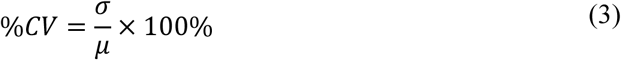

Here, μ and σ represent the mean and standard deviation of the positive (p) and negative (n) droplet fluorescence amplitudes, respectively. Performance metrics were calculated from reaction-level summary statistics and averaged between technical replicates.

### RPA primer design and screening

Candidate RPA primers targeting the *dnaJ* gene were designed using Geneious Prime 2025.2.2 (https://www.geneious.com) based on a consensus sequence (Text S1) generated from 14 *V. coralliilyticus dnaJ* sequences retrieved from GenBank (Table S1). Primer design followed the manufacturer’s recommended guidelines for RPA assays (TwistDx, United Kingdom). Candidate primers were evaluated *in silico* using Integrated DNA Technologies’ (IDT) OligoAnalyzer™ Tool (Owczarzy et al., 2008) to assess primer characteristics, including melting temperature and potential secondary structures. Primer specificity was further evaluated using NCBI BLASTN and NCBI Primer-BLAST (Benson et al., 2013). Seventeen primer candidates resulting in 26 forward/reverse combinations and targeting two overlapping regions on the *dnaJ* gene were selected for experimental screening (Table S2). Primers were synthesized commercially (Merck Group, Germany).

RPA reactions were performed using the TwistAmp Basic Kit (TwistDx), according to the manufacturer’s instructions. Reactions were incubated at 40 °C for 20 min. After incubation, reactions were heated to 95 °C for 5 min to terminate amplification and standardize reactive endpoints. Amplified products were purified using the Isolate II PCR and Gel Kit (Meridian Bioscience, USA), separated by electrophoresis on 2% agarose gels, imaged under UV transillumination, and qualitatively assessed for target amplification and nonspecific products.

Primer screening was initially performed using genomic DNA from the *V. coralliilyticus* type-strain (0.1 ng µL^−1^) to evaluate amplification across candidate primer pairs. Preliminary screening identified a single target region that produced the most specific amplification and was therefore selected for further primer evaluation. All primer pairs within this region were assessed for off-target amplification using genomic DNA from seven non-target *Vibrio* species (0.1 ng µL^−1^; Table 1), with a no-template control (NTC) included in each reaction strip.

### crRNA design, screening, and assay optimization

Candidate crRNAs were designed in Geneious Prime 2025.2.2 using the d*naJ* consensus sequence. Target sites adjacent to the Cas12a 5’-TTTV-3’ protospacer adjacent motif were identified in the *dnaJ g*ene. Candidate crRNAs were evaluated *in silico* for target specificity using NCBI BLASTN and four candidate crRNAs were selected for experimental validation (Table S3).

CRISPR detection reactions used a custom fluorophore-quenched ssDNA reporter (5’-FAM-TTATT-BHQ-3’), Alt-R™ *Lachnospiraceae bacterium* Cas12a (Cpf1) Ultra, and custom Alt-R™ *L.b.* Cas12a crRNAs (IDT; USA). Cas12a/crRNA ribonucleoprotein complexes were pre-assembled by incubating Cas12a and crRNA (1:1) at 37°C for 15 min in a buffer containing 20 mM Tris-HCl (pH 7.5) and 150 mM KCl. Detection reactions (20 μL) comprised 50 nM Cas12a and 50 nM crRNA pre-assembled, 500 nM reporter, reaction buffer (20 mM Tris-HCl pH 7.5; 150 mM KCl; 7.5 mM MgCl2), and 2 μL RPA product. Fluorescence was monitored at 37°C for up to 15 min using a Genie II Mk2 fluorometer (GeneWorks, Australia). Endpoint fluorescence measured after 10 min was used for all comparative analyses, with reactions exceeding 5,000 relative fluorescence units (RFU) classified as positive. This conservative threshold was chosen to minimize the likelihood of false-positive interpretation according to initial runs of positive and negative control samples.

crRNA performance was initially evaluated using genomic DNA from the *V. coralliilyticus* type-strain (0.1 ng µL⁻^1^), after which the best performing crRNAs were assessed against DNA of all seven *V. coralliilyticus* strains (0.1 ng µL⁻^1^) with tests in triplicate. The effect of RPA incubation time (10 verses 20 min) on assay performance was evaluated both at the RPA stage and at the CRISPR detection stage. Amplification specificity was assessed using *V. coralliilyticus* type-strain DNA (0.1 ng µL⁻^1^), an off-target DNA mixture (OT mix) comprising equal concentrations of the seven non-target *Vibrio* species DNA (0.1 ng µL⁻^1^), OT mix without *Vibrio brasiliensis*, and NTCs. The exclusion of *V. brasiliensis* was based on initial explorative runs per off-target species using the same condition. RPA products were cleaned using the Isolate II PCR and Gel Kit, separated on 2% agarose gel, and imaged with UV transillumination. CRISPR detection specificity was assessed using *V. coralliilyticus* type-strain DNA (0.1 ng µL⁻^1^), OT mix, and NTCs with six replicates reactions per treatment. Pairwise comparisons among templates were performed using estimated marginal means with Holm-adjusted p-values.

### Sensitivity assessment

Analytical sensitivity was evaluated using a serial dilution of *V. coralliilyticus* type-strain genomic DNA spanning 0-774 *dnaJ* target copies of per reaction, with target concentrations determined by ddPCR. Each dilution was tested using the developed RPA-CRISPR/Cas12a assay alongside an NTC, with eight technical replicates per test concentration. The limit of detection (LOD) was defined as the lowest target concentration that yielded positive detection in 100% of replicate reactions, in accordance with the criteria described by De Brauwer et al. (2022).

### Evaluation using aquaculture water samples

Water samples were collected from two 1,150 L flow-through aquaria (2,800 x 900 x 450 mm) holding *Acropora tersa* broodstock corals at the Australian Institute of Marine Science. Coral samples were collected under the Great Barrier Reef Marine Park Zoning Plan 2003, Section 5.4 Authorization for the Pilot Deployments Program. One tank contained six colonies exhibiting active tissue necrosis (diseased tank), while the second contained 15 visually healthy colonies (healthy tank). In both systems, colonies were positioned on racks evenly distributed throughout the tank at mid-water depth (approximately 20 cm). Within each tank, four replicate 50 mL water samples were collected from three locations (Figure 1A): inflow water, water adjacent to coral colonies (3-5 cm), and water near the tank outflow approximately 1 m from the nearest coral colonies (hereafter referred to as the “inflow”, “close”, and “far” sampling locations, respectively). Water was collected using sterile 50 mL syringes and immediately filtered through sterile 0.22 µm polyether sulfone membranes mounted in sterile Swinnex filter housings. Filters were stored in 432 µL Qiagen Buffer ATL (Qiagen, Germany) until DNA extraction using a modified Maxwell RSC PureFood GMO and Authentication Kit protocol (Text S2) on a Maxwell RSC Instrument (Promega, USA).

**Figure 1.**
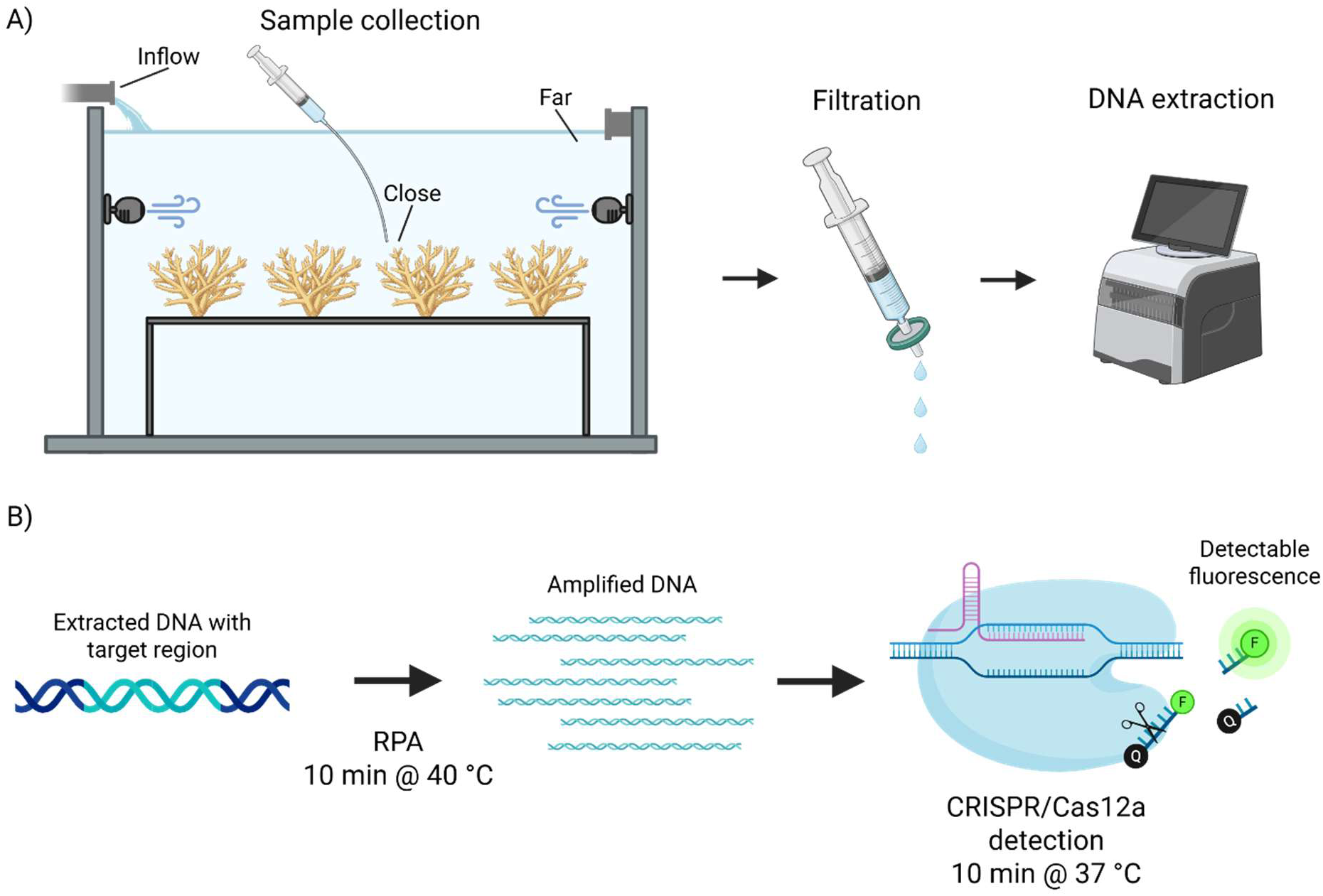
Aquaculture water test and workflow for the rapid detection of *Vibrio coralliilyticus*. (A) Sample collection and processing for the aquaculture water test. (B) RPA-CRISPR/Cas12a assay workflow. The CRISPR detection produces a fluorescent signal when bound to amplified target DNA, wherein Cas12a collaterally cleaves reporter molecules, separating the quenchers (Q) from the fluorophores (F). Figure created in BioRender (https://BioRender.com/ihkrxju).

Purified DNA was analyzed using both the developed RPA-CRISPR/Cas12a assay and ddPCR for target detection and quantitative benchmarking. For the RPA-CRISPR/Cas12a assay, 5 µL of template DNA was added to each reaction, whereas 2 µL was used for each ddPCR reaction to avoid positive droplet saturation. All samples were analyzed in duplicate technical replicates, with two positive-template controls and two no-template controls (NTCs) included in each assay run to verify assay performance and rule out potential contamination. To assess differences in target concentrations between sampling locations, mean ddPCR target copy numbers were compared between close and far samples within diseased and healthy tanks using a two-factor ANOVA. Pairwise comparisons were adjusted using the Holm method to account for multiple testing. Estimated marginal means were additionally used to compare mean target concentrations between healthy and diseased samples.

## Results

### ddPCR assay optimization

Primer concentration and annealing temperature had a marked influence on ddPCR assay performance (Figure S1). Across the primer concentration gradient, the highest overall assay performance was achieved using 200 nM primers, which produced the greatest separation between positive and negative droplet populations (mean Z′ = 0.77; mean SNR = 2.77) while maintaining low fluorescence variability within positive droplets (mean %CV = 3.6). Increasing primer concentration progressively reduced droplet cluster separation, with 700 nM yielding the lowest mean Z′ (0.67) and SNR (1.92), although fluorescence variability remained low across all primer combinations (%CV generally < 5). Optimal performance was observed at an annealing temperature of 52 °C and a primer concentration of 200 nM (Z′ = 0.82; SNR = 2.97; %CV of positive droplets = 3.1), and thus this primer concentration was used in all subsequent analyses.

Annealing temperature strongly influenced assay specificity (Table S4). Non-target amplification declined with increasing temperatures and was largely eliminated at temperatures ≥ 63.3 °C, where all seven on-target *Vibrio* species with non-target samples yielding ≤ 0.03 copies µL^−1^. However, target amplification also decreased at these higher temperatures, with Z′ values falling below 0.54 for two of five *V. coralliilyticus* strains tested at 63.3 °C. In contrast, an annealing temperature of 60.7 °C maintained robust amplification across all five target strains (Z′ = 0.82-0.85; SNR = 3.97-4.22; %CV of positive droplets = 2.8 −3.3), while all seven non-target species remained effectively negative (< 0.09 copies µL^−1^). At temperatures ≤ 57.5 °C, non-specific amplification increased among non-target species. Consequently, 60°C was selected as the ddPCR annealing temperature for all subsequent experiments as the optimal compromise between analytical sensitivity and assay specificity.

### RPA primer screening

All 26 RPA primer combinations successfully amplified *V. coralliilyticus* type-strain DNA; however, varying levels of off-target amplification were observed among the seven non-target *Vibrio* species (Table S5). Primer pair 412F (5’-TCA AAA GAA ATT GAA GTT CCA ACG CTA GTT AA-3’) and 620R (5’-CCA TGA CAT GAG TTA CAA GGA TCT TTG ATG AT-3’) exhibited the highest specificity across all primer pair candidates, producing the lowest level of off-target amplification. The 20 min RPA incubation produced a faint band with the OT mix template, but not with the OT mix without *V. brasiliensis* (Figure 2), supporting primer screening results that *V. brasiliensis* was the sole non-target species which amplified with the 412F/620R primer pair. This non-target amplification was not observed with the 10 min incubation. Although primer combination 412F/619R displayed a similar specificity profile (Table S5), it generated a more intense target-sized amplicon from *V. brasiliensis* DNA and was therefore not selected.

**Figure 2.**
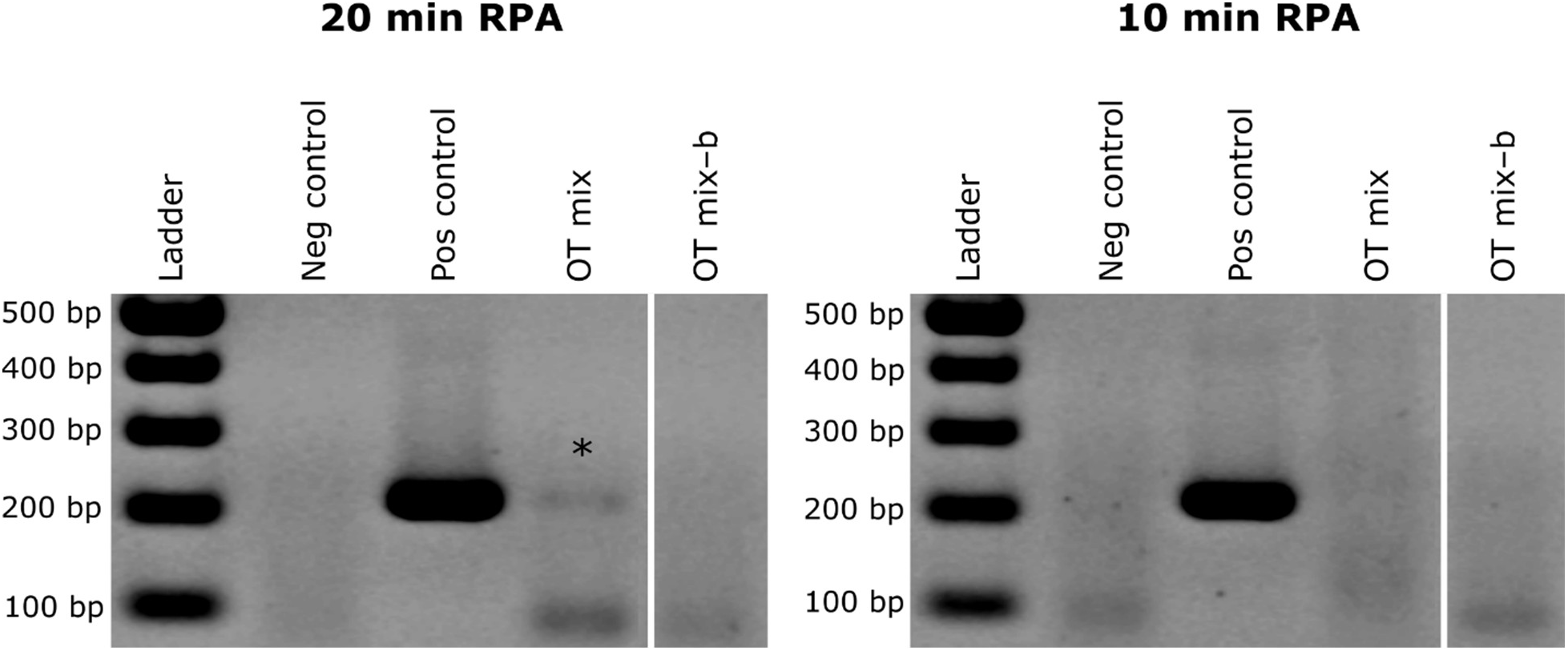
Effect of incubation length on recombinase polymerase amplification (RPA) specificity. Visualized with 2% agarose gel electrophoresis. Wells include 100 bp DNA ladder, no-template control (neg control), *Vibrio coralliilyticus* type-strain (pos control), a mix containing DNA of seven non-target *Vibrio* spp. (OT mix), and OT mix without *V. brasiliensis* (OT mix–b). Target size is 209 bp. Asterisk indicates off-target amplification (note that the off-target RPA amplicons did not affect CRISPR/Cas12 signals; see below). See Table 1 for list of bacterial isolate DNA used. These images have been uniformly transformed (contrast and aspect ratio) for better visibility (see Figure S2 for the unaltered images).

### crRNA screening

The four candidate crRNAs differed markedly in their ability to detect *V. coralliilyticus* type-strain DNA (Figure 3). After 10 min of incubation, crRNA_1 and crRNA_3 generated substantially higher fluorescence signals than crRNA_2 and crRNA_4 (Figure 3A), which were therefore excluded from further evaluation. crRNA_1 and crRNA_3 were subsequently assessed for cross-strain target detection. Cross-strain screening revealed that crRNA_1 produced consistently strong fluorescence signals across all *V. coralliilyticus* strains tested, with the exception of strain C2 (Figure 3B). In contrast, crRNA_3 exhibited greater variability among strains and generally yielded lower fluorescence signals than crRNA_1. The only exception was strain C2, for which crRNA_3 generated approximately fivefold higher fluorescence than crRNA_1. Given its superior and more consistent performance across Indo-Pacific isolates, crRNA_1 (5’-[TTTG] TGA TGG CAG TGG CGC GAA GAA-3’) was selected for all subsequent experiments. Although crRNA_3 showed improved detection of the Caribbean isolate C2, this strain was outside the primary geographical scope of the assay.

**Figure 3.**
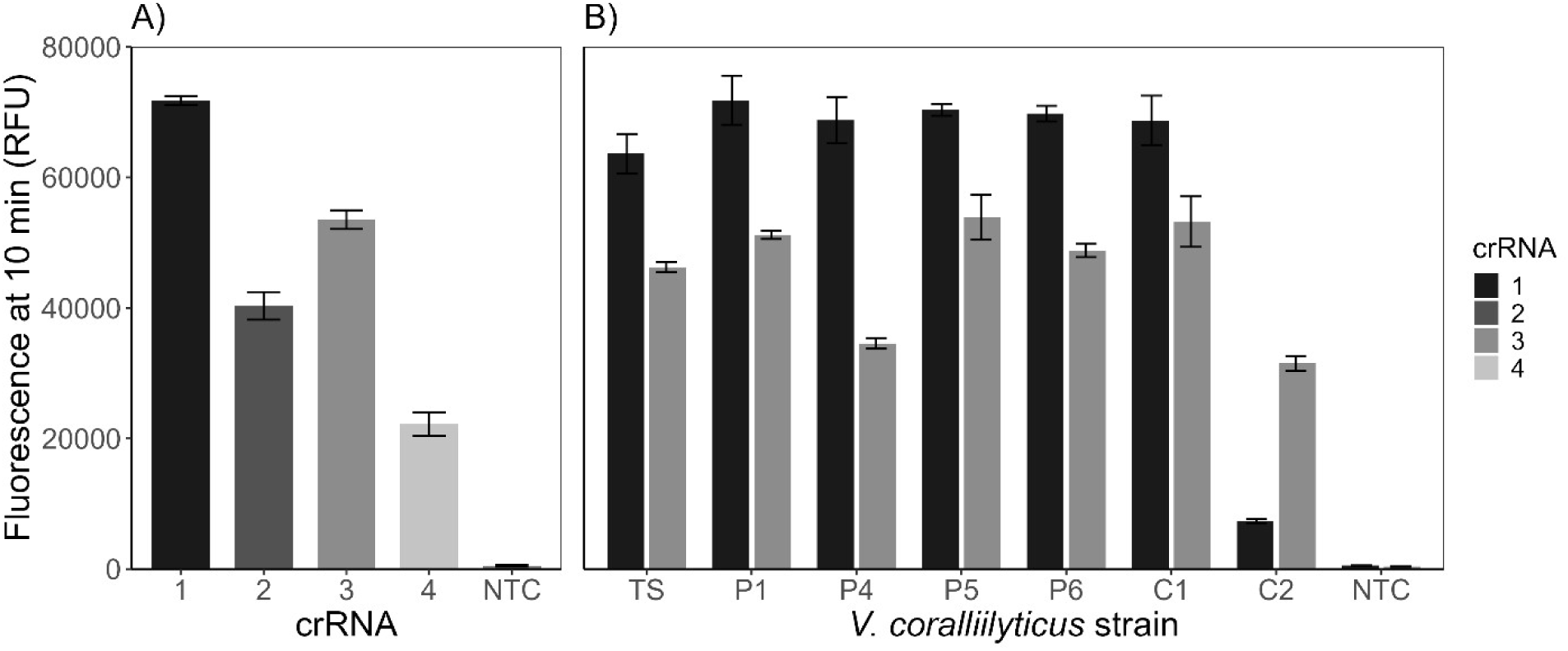
CRISPR-RNA (crRNA) screening results. Fluorescence production in RPA-CRISPR/Cas12a detection reactions targeting *Vibrio coralliilyticus*. Fluorescence is reported in relative fluorescence units (RFU) after 10 min CRISPR incubation. (A) Fluorescence production observed across four crRNA candidates with template inputs of *V. coralliilyticus* type-strain and no-template controls (NTC). (B) Fluorescence production observed across seven different strains of *V. coralliilyticus* (Table 1) and NTCs utilizing crRNA_1 (black) and crRNA_3 (medium gray). Error bars depict standard error.

### Effect of RPA incubation time on CRISPR/Cas12a signals

Despite the RPA amplification in the OT-mix depending on the incubation time, the RPA incubation time (20 versus 10 min) had no detectable effect on CRISPR/Cas12a fluorescence output or assay specificity. Mean fluorescence generated from *V. coralliilyticus* type-strain DNA was comparable following 20 min (59,109 ± 2,742 RFU) and 10 min (59,952 ± 1,915 RFU) RPA amplification, with no significant difference between incubation times (p = 0.688). Similarly, fluorescence from the OT-mix including *V. brasiliensis* (98 ± 106 vs. 111 ± 105 RFU; p = 0.995) and NTCs (127 ± 160 vs. 87 ± 157 RFU, p = 0.988) remained consistently low under both conditions. As no reduction in assay performance was observed, a 10 min RPA incubation was adopted for all subsequent experiments, reducing the total assay time without compromising sensitivity or specificity.

### Sensitivity assessment

The analytical sensitivity of the RPA-CRISPR/Cas12a assay was determined using a ddPCR-quantified dilution series of *V. coralliilyticus* DNA (Figure 4). Positive detection in all replicates containing ≥ 9 *dnaJ* target copies per reaction was observed, whereas the detection rate declined to 62.5% at 3 copies per reaction and decreased further at lower target concentrations. Accordingly, the LOD was established at 9 *dnaJ* target copies per reaction. Endpoint fluorescence remained high across concentrations from 76 to 9 target copies per reaction, ranging from 80,400 ± 4,517 RFU to 58,109 ± 14,662 RFU, before declining sharply at lower template concentrations. These results demonstrate that the assay maintained robust fluorescence output and reliable detection close to its analytical detection limit.

**Figure 4.**
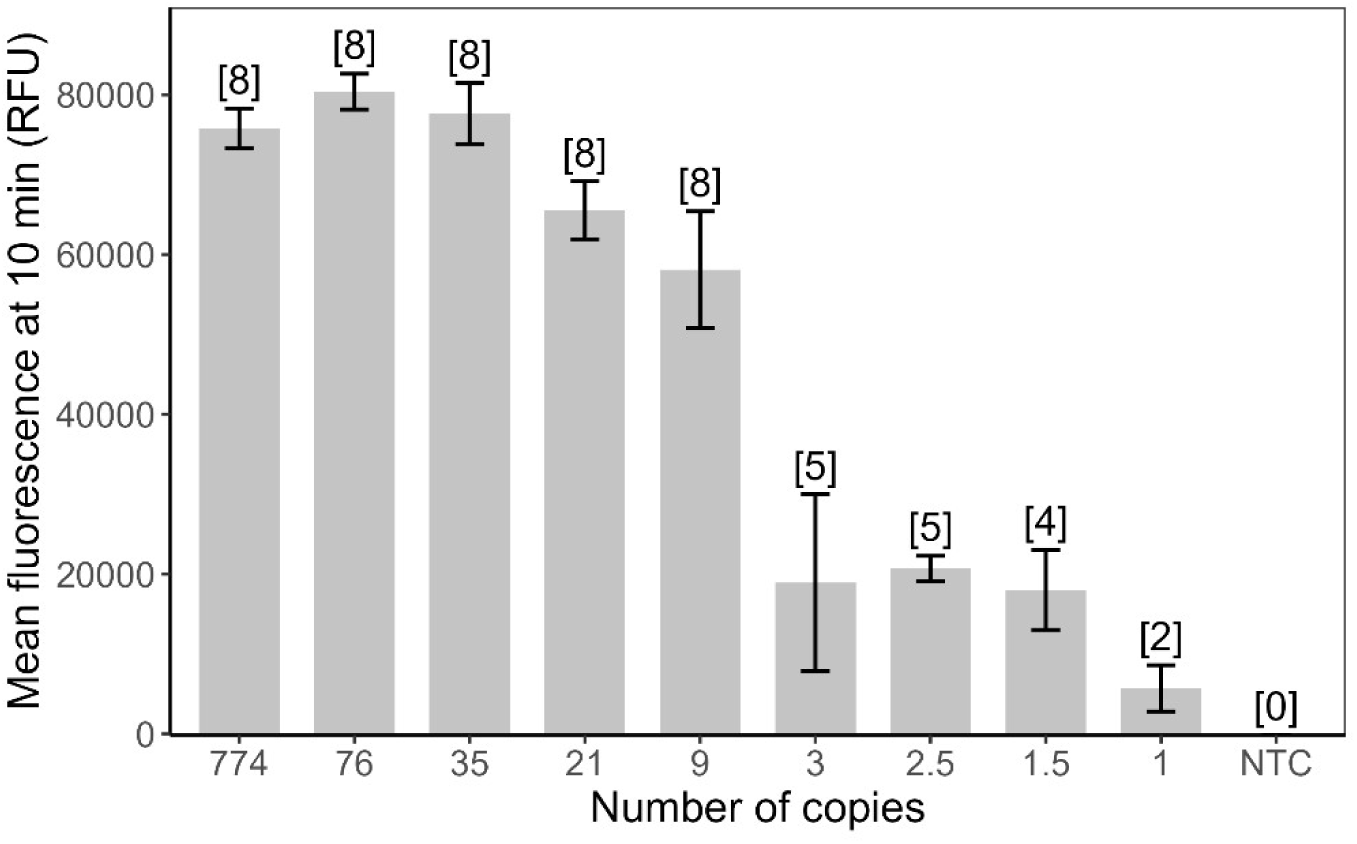
Assay sensitivity assessment results. Mean fluorescence generated with each dilution after 10 min CRISPR incubation. Number of target DNA copies in each dilution and no-template control (NTC) were established by droplet digital PCR. Numbers in brackets above each bar correspond to technical replicates out of 8 for that dilution which produced detection with the RPA-CRISPR/Cas12a assay. Fluorescence is reported in relative fluorescence units (RFU). Error bars depict standard error.

### Evaluation using aquarium water samples

The RPA-CRISPR/Cas12a assay detected *V. coralliilyticus* in 100% of water samples collected from tanks holding both diseased and healthy *Acropora tersa* colonies, whereas all inflow water or NTC samples were negative (Table 2). Results were consistent with ddPCR analyses, which confirmed the presence of *V. coralliilyticus* in all coral-containing tanks, but ddPCR quantification showed considerably higher *V. coralliilyticus* concentrations in diseased tanks than in tanks holding healthy corals (p = 0.0003). Within diseased tanks, mean target abundance in samples collected adjacent to coral colonies was approximately double that of samples collected near the outflow, with this difference reaching marginal significance (p = 0.0497). In contrast, no spatial differences were observed in tanks holding healthy colonies (p = 0.9719), where target abundance remained consistently low (<1,000 *dnaJ* copies per reaction). Inflow water and NTC samples showed negligible or undetectable target DNA, consistent with the absence of RPA-CRISPR/Cas12a detections.

**Table 2.** Results from the aquaculture water test of the RPA-CRISPR/Cas12a assay cross-validated with ddPCR quantification. Sample groups (n = 4 replicates sample^−1^) included water taken adjacent to the corals (Close), near the tank outflow (Far), from the inflow water, and no-template controls (NTC). RPA-CRISPR/Cas12a detection results are given as detection rate of *V. coralliilyticus* by sample group. ddPCR results quantifying *V. coralliilyticus* concentration are reported as the average *dnaJ* copy number equivalent to what was present in each RPA-CRISPR/Cas12a detection reaction.

| Treatment | Sample group | RPA-CRISPR/Cas12a detection rate (%) | Mean copies reaction<sup>-1</sup> | SE | Close vs. Far p-value |
| --- | --- | --- | --- | --- | --- |
| **Diseased** | Close | 100 | 18,211.98 | 2,707.56 | 0.0497 |
|  | Far | 100 | 10,015.27 | 2,232.7 |  |
|  | Inflow | 0 | 0.79 | 0.52 |  |
| **Healthy** | Close | 100 | 696.32 | 175.73 | 0.9719 |
|  | Far | 100 | 871.1 | 60 |  |
|  | Inflow | 0 | 0.82 | 0.53 |  |
|  | NTC | 0 | 0 | 0 |  |

*Note.* SE = Standard error. Close-Far comparisons were performed using mean ddPCR concentrations from four replicates per sample group, with p-values adjusted using the Holm method.

## Discussion

### Development of a rapid, sensitive, and specific diagnostic assay

The RPA-CRISPR/Cas12a workflow developed here (see Figure 1B) enabled detection of *V. coralliilyticus* in less than 30 min after DNA extraction, including reaction preparation and transfer of amplification products. Following optimization, the assay comprised 10 min of RPA amplification followed by 10 min of CRISPR/Cas12a mediated detection, demonstrating the potential of this approach for rapid pathogen screening. See Text S3 for a succinct protocol of the developed assay, including probe sequences and reagent concentrations. The ability to achieve rapid and specific detection is particularly relevant for marine *Vibrio* diagnostics, where high genetic similarity among closely related species can make development of species-specific molecular assays challenging (Leunda-Esnaola et al., 2024; Thompson et al., 2005).

Extensive off-target screening during primer optimization was important for achieving the specificity required for environmental application. Several primer combinations that efficiently amplified *V. coralliilyticus* also produced amplification with closely related *Vibrio* species. RPA operates under less stringent amplification conditions than conventional PCR and can therefore be more tolerant of greater primer-template mismatches, increasing the potential for non-specific amplification (Tan et al., 2022; Higgins et al., 2022). The extensive screening here against related marine *Vibrio* spp. therefore represents an important component of assay development and provides greater confidence in the specificity of the selected primers than would be achieved through minimal non-target screening reported in previous published *Vibrio* RPA assays (Dong et al., 2020; Geng et al., 2019; X. Yang et al., 2021; Zhong et al., 2025; Zhu et al., 2021). Notably, initial RPA primer screening was conducted using a 20 min RPA incubation, longer than the 10 min amplification period adopted in the final assay. This extended incubation period provided a deliberately stringent test of primer specificity because prolonged amplification increases the opportunity for non-specific products to accumulate. Primers that remained specific under these conditions were therefore considered better suited to subsequent application in environmentally complex coral aquaculture samples.

Despite the non-specific RPA amplification seen with the longer incubation time, our results demonstrated that the design of the crRNA provides an additional layer of target specificity and can prevent these unspecific products from producing the final CRISPR/Cas12a signals. Screening of the crRNAs further highlighted the importance of accounting for strain-level genetic variation during assay development. crRNA_1 generated consistently strong fluorescence across most *V. coralliilyticus* strains tested, although strain C2 produced substantially weaker fluorescence relative to the other strains. Detection inconsistencies involving strain C2 have also been reported in previous work targeting *dnaJ* (Pollock et al., 2010), suggesting that sequence variation within or surrounding the target region may influence assay performance. Nevertheless, crRNA_1 provided robust detection across strains more representative of Indo-Pacific coral systems and was therefore selected for the final assay. Where detection of strain C2 is a specific priority, crRNA_3 may provide a suitable alternative. Strong fluorescence generated by crRNA_1 is also advantageous for future deployment using simplified visualization approaches, including blue-light fluorescence detection, which may reduce the need for dedicated laboratory instrumentation (Yang et al., 2025).

### Performance as a coral aquaculture monitoring tool

The assay successfully detected *V. coralliilyticus* DNA in water collected from coral aquaculture tanks containing both diseased and visually healthy *Acropora tersa* colonies, but not in inflow water. Presence-absence detection patterns in the RPA-CRISPR/Cas12a assays were consistent with ddPCR quantifications, while marked difference in abundance based on the tank treatment and sampling locations were unraveled by ddPCR quantification. The absence of amplification in inflow water and no template control samples further supports the specificity of the RPA-CRISPR/Cas12a assay under these conditions and provides no evidence of cross-contamination during the experiment. Collectively, these findings demonstrate that the RPA-CRISPR/Cas12a assay reliably detected the presence of *V. coralliilyticus* in aquaculture water samples, while ddPCR provided quantitative resolution needed to distinguish low background levels from elevated pathogen loads associated with diseased coral colonies.

Importantly, detection was achieved directly from tank water DNA rather than DNA extracted from purified cultures, demonstrating that the assay can function in a biologically complex sample matrix. This represents an important step towards environmental applications because molecular assays frequently perform differently on environmental samples due to increased biological and chemical complexity. For example, aquaculture water samples contains diverse non-target DNA and potentially inhibitory compounds that can reduce amplification efficiency and, consequently, affect downstream CRISPR-based detection performance (Gentry-Shields et al., 2013). The successful detection of *V. coralliilyticus* in these samples indicates that the assay retains sufficient analytical performance despite this additional complexity. However, validation across a broader range of aquaculture systems, water chemistries, colony conditions, and naturally occurring pathogen concentrations will be necessary before the assay can be considered operationally ready.

The rapid turnaround time of the assay offers a potential advantage over conventional laboratory diagnostics such as ddPCR and qPCR, particularly in coral aquaculture where disease outbreaks occur rapidly and biomass losses can be substantial (Sheridan et al., 2013). A diagnostic capable of producing results within approximately 30 min could support more responsive health surveillance, allowing managers to increase monitoring, isolate potentially affected colonies, or initiate confirmatory quantitative testing before disease progression becomes widespread. Importantly, the intended value of such an assay is not necessarily to replace quantitative laboratory diagnostics, but to provide a rapid screening tool that can inform when more intensive investigation is warranted.

Future simplification of the assay will also be important for practical deployment. In particular, integrating RPA and CRISPR reactions into a single reaction tube could reduce handling steps, shorten the workflow, and minimize the risk of aerosolized cross-contamination (Li et al., 2025). Reducing instrumentation requirements represents a complementary avenue for improving accessibility. Preliminary testing in the present study demonstrated that positive reactions could be visually distinguished using an inexpensive blue-light fluorescence viewer (P51™ Molecular Fluorescence Viewer; miniPCR bio, USA), suggesting that dedicated fluorescence instrumentation may not be essential for qualitative detection (Figure S3). This approach requires further validation, particularly to establish detection limits and the reliability of visual interpretation but highlights the potential to advance the assay from a laboratory proof-of-concept toward a practical diagnostic capable of rapid health surveillance in coral aquaculture.

Interpretation of a binary presence/absence detection result is ecologically complex, since detection of *V. coralliilyticus* does not necessarily indicate active disease because the bacterium may occur at a low abundance within healthy coral holobionts (Munn, 2015). Consistent with this, ddPCR quantification in the present study detected low numbers of *V. coralliilyticus* in samples from tanks holding healthy corals. Consequently, a highly sensitive presence/absence assay could potentially identify low-level background populations without indicating an immediate disease threat. The practical value of rapid detection will therefore depend not only on analytical sensitivity and specificity, but also on understanding how pathogen abundance relates to host condition and disease progression.

Corroborating with our study, evidence from other aquaculture systems suggests that pathogen load can provide more informative estimates of disease risk than pathogen presence alone. For bacterial and viral diseases of crustacean aquaculture species, pathogen load is commonly associated with infection severity and outbreak risk, and qPCR-based thresholds have been incorporated into health management frameworks (Durand & Lightner, 2002; Zhang et al., 2025). Equivalent pathogen load-disease relationships remain comparatively poorly characterized in corals, although available evidence suggests that increasing *V. coralliilyticus* abundance may be associated with declining host condition. For example, heat-stressed *Pocillopora damicornis* exhibited a four-order-of-magnitude increase in *V. coralliilyticus* abundance alongside microbial dysbiosis and declining health (Tout et al., 2015). The next stage of diagnostic development should move beyond simple presence/absence pathogen detection towards approaches capable of distinguishing background colonization from pathogen loads associated with elevated disease risk. The modularity of CRISPR-based diagnostics provides a useful platform for this development because target recognition, amplification, and signal generation can be independently modified (Kaminski et al., 2021). Rather than attempting to replace quantitative laboratory approaches, future development of this assay could focus on establishing rapid quantitative or semi-quantifying information sufficient to distinguish baseline from elevated pathogen loads. Such an approach may offer a useful compromise between the speed and accessibility of field-based RPA-CRISPR assays and the quantitative information provided by qPCR or ddPCR approaches. One promising strategy is to constrain the amplification period such that fluorescence signals remain dependent on initial target template concentration rather than reaching a saturated endpoint. This concept was recently demonstrated using an RPA-CRISPR/Cas12a assay for cytomegalovirus, where shortened RPA amplification enabled quantitative discrimination relative to PCR-based measurements (Chao et al., 2026). Applying a similar strategy to *V. coralliilyticus* could potentially enable rapid categorization of pathogen loads without requiring the instrumentation and processing time associated with conventional quantitative assays.

Further improvements could also be achieved by targeting biomarkers more directly associated with virulence activity or disease progression, rather than organism presence alone. For example, virulence-associated genes such as *vcpA* may provide greater ecological relevance for assessing disease risk (Ushijima et al., 2020; Wilson et al., 2013). Together these approaches provide a pathway towards diagnostics that are not simply pathogen specific, but more closely aligned with the biological processes underlying coral disease.

## Conclusion

This study demonstrates the feasibility of RPA-CRISPR/Cas12a as a rapid and specific approach for detecting *V. coralliilyticus* in coral aquaculture water samples. The assay combined short amplification and detection times with robust specificity across tested *V. coralliilyticus* strains and successful detection from environmentally derived aquaculture water. These characteristics highlight the potential of RPA-CRISPR/Cas12a diagnostics as a rapid screening tool for coral health surveillance. Importantly, pathogen detection alone is unlikely to provide a complete measure of disease risk, where *V. coralliilyticus* can occur at low abundance in apparently healthy corals. Future development should therefore focus on rapid semi-quantitative detection, biologically informative targets, simplified visualization, and integration of amplification and CRISPR detection into a one-pot format to achieve a point-of-need implementation. As coral aquaculture activities continue to expand, rapid diagnostic assays of this design could complement quantitative laboratory approaches and support proactive health management in coral aquaculture by enabling frequent surveillance and earlier interventions following pathogen detection.

## Author Contributions

Study design and conception: YS, JD. Assay design and development: SA, YS, JD. Acquisition, analysis, and interpretation of data: SA, JD, CMG, YS. Writing of original draft: SA. Writing review & editing: SA, JD, CMG, DGB, YS.

## Data Availability Statement

All bacterial sequences generated in this research were uploaded to NCBI GenBank under accession numbers PZ785359 - PZ785372 listed in Table 1. Assay probe, consensus sequence, and ddPCR optimization data are available in the supplemental materials.

## Acknowledgements

This research was undertaken within the Reef Restoration and Adaptation Program (RRAP), funded by the partnership between the Australian Government’s Reef Trust, the Great Barrier Reef Foundation, and research partners including the Australian Institute of Marine Science and James Cook University. We acknowledge the Bindal and Wulgurukaba peoples as the Traditional Owners and Custodians of the land and Sea Country in which this work was conducted. We also gratefully appreciate the anonymous reviewer input to this publication.

## Conflicts of Interest

The authors report no conflicts of interest.

---

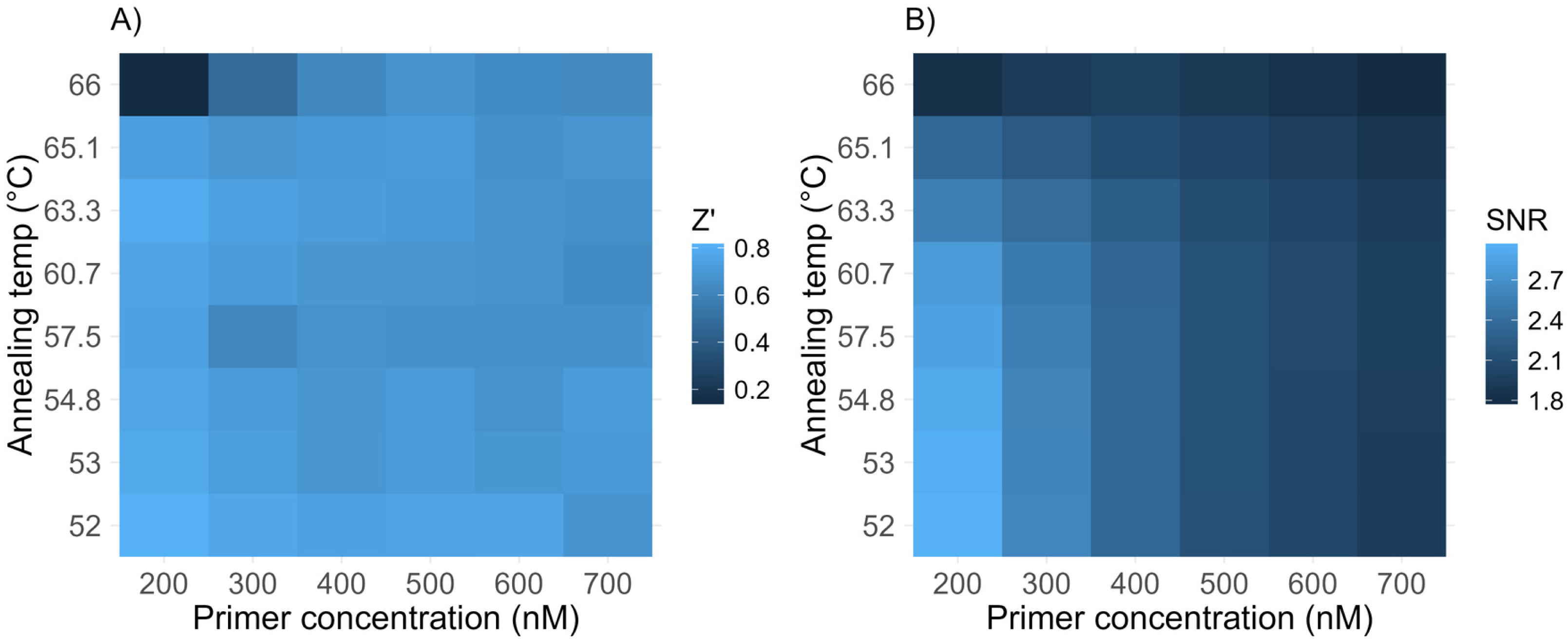

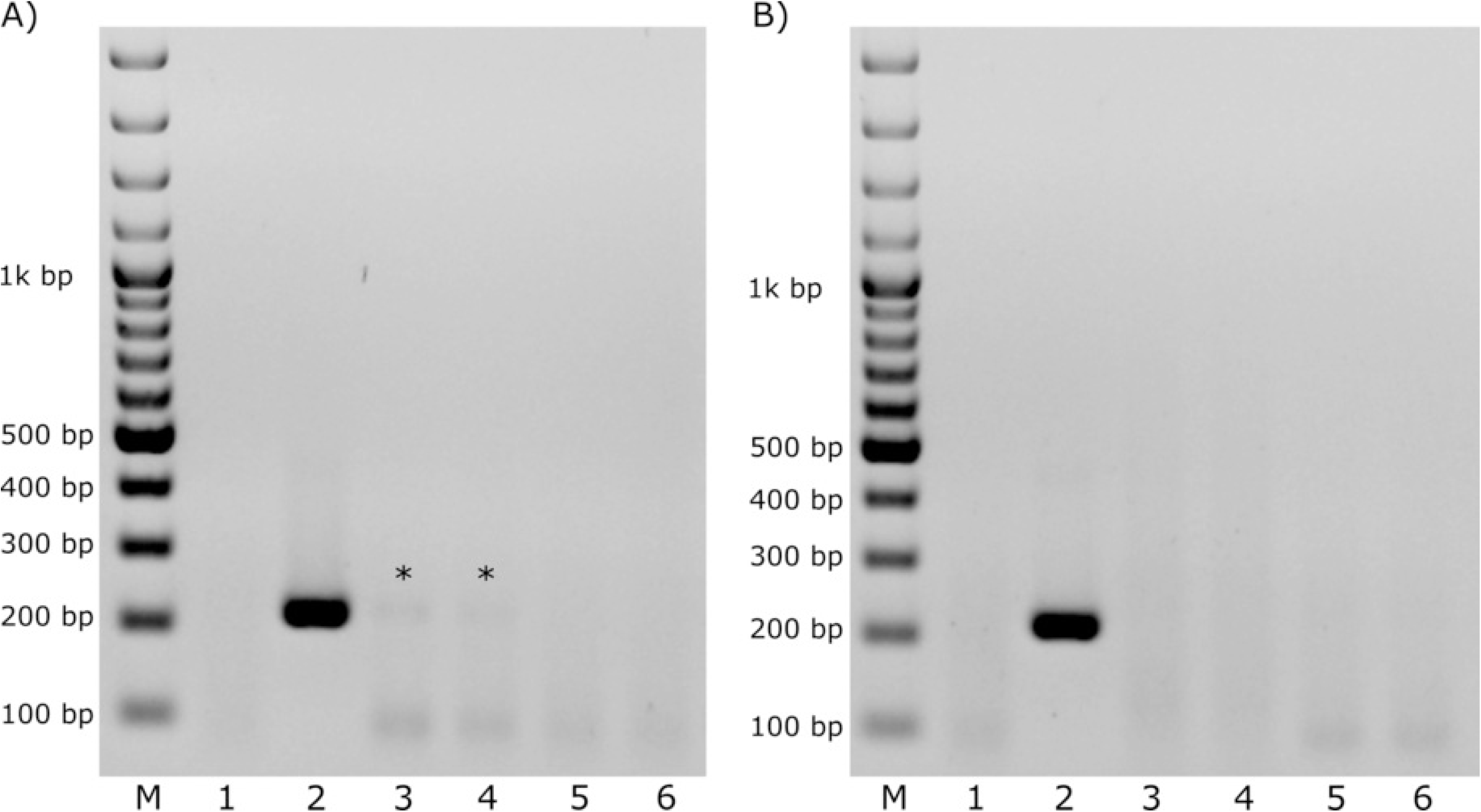

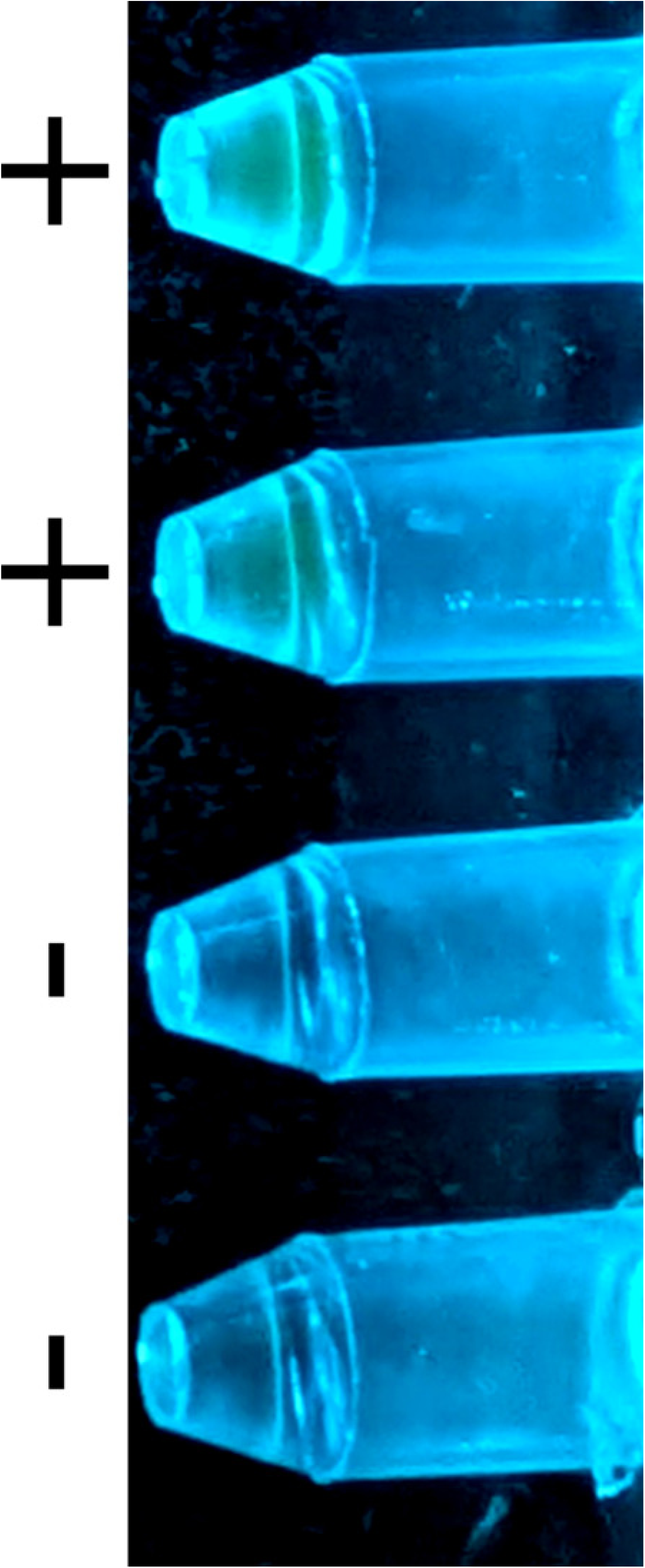

